# Post-decision wagering after perceptual judgments reveals bi-directional certainty readouts

**DOI:** 10.1101/272872

**Authors:** Caio M. Moreira, Max Rollwage, Kristin Kaduk, Melanie Wilke, Igor Kagan

## Abstract

Humans and other animals constantly evaluate their decisions in order to learn and behave adaptively. Experimentally, such evaluation processes are accessed using metacognitive reports made after decisions, typically using verbally formulated confidence scales. When subjects report high confidence, it reflects a high certainty of being correct, but a low confidence might signify either low certainty about the outcome, or a high certainty of being incorrect. Hence, metacognitive reports might reflect not only different levels of decision certainty, but also two certainty directions (certainty of being correct and certainty of being incorrect). It is important to test if such bi-directional processing can be measured because, for decision-making under uncertainty, information about being incorrect is as important as information about being correct for guidance of subsequent behavior. We were able to capture implicit bi-directional certainty readouts by asking subjects to bet money on their perceptual decision accuracy using a six-grade wager scale (post-decision wagering, PDW). To isolate trial-specific aspects of metacognitive judgments, we used pre-decision wagering (wagering before the perceptual decision) to subtract, from PDW trials, influences resulting from non-trial-specific assessment of expected difficulty and psychological biases. This novel design allowed independent quantification of certainty of being correct and certainty of being incorrect, showing that subjects were able to read out certainty in a bi-directional manner. Certainty readouts about being incorrect were particularly associated with metacognitive sensitivity exceeding perceptual sensitivity (i.e. meta-d′ > d′), suggesting that such enhanced metacognitive efficiency is driven by information about incorrect decisions. Readouts of certainty in both directions increased on easier trials, and both certainty directions were also associated with faster metacognitive reaction times, indicating that certainty of being incorrect was not confounded with low certainty. Finally, both readouts influenced the amount of money subjects earned through PDW, suggesting that bi-directional readouts are important for planning future actions when feedback about previous decisions is unavailable.

## 1. Introduction

Humans and other animals are able to assess their own cognitive processes (perception, memory and decisions) to flexibly adapt their behavior (Fleming and Lau, 2014; Hampton, 2009; Kepecs and Mainen, 2012). This metacognitive assessment can be understood as readouts of a varying certainty – probability distributions over contributing random variables – associated with inherently uncertain sensory evidence and cognitive processes (Kepecs, 2013; Ma and Jazayeri, 2014; Pouget et al., 2016) and is especially useful for planning post-decisional actions under uncertainty (Fleming et al., 2012a; Kepecs and Mainen, 2012; Kiani et al., 2014).

Most previous work on assessing decision certainty utilized confidence judgments (Fleming et al., 2012; Hebart et al., 2014; Heereman et al., 2015; Persaud et al., 2007), which can be computationally defined as subjective probability of having done a correct decision (Pouget et al., 2016). However, while confidence is a form of certainty about being correct, these measures are not equivalent (Fleming and Daw, 2017; Pouget et al., 2016). For example, a recent study indicated that there might be a continuum in the knowledge one has about having done something wrong and something right (Boldt & Yeung, 2015) and, in this context, confidence might vary with certainty levels but also with what we name *‘ certainty directions’* : one direction for the *certainty of being correct*, and the opposite direction for the *certainty of being incorrect*. Hence, confidence ranges from 0 (high certainty of being incorrect) to 1 (high certainty of being correct) with confidence around 0.5 corresponding to intermediate levels of certainty of either being correct or incorrect. Such bi-directional certainty might influence future actions in a continuous bi-directional way. Imagine a man in a hurry going to the supermarket and deciding whether to enter an aisle to search for a specific product. If he is uncertain about his decision (confidence around 0.5), he will probably slow down and search for the product from a distance, staying close to other aisle options. This might not be the best way to find the product, but it is the best way to avoid spending the main resource (in this case, time) on this uncertain decision. If he is highly certain about the decision, the planning of a next action might have two distinct outcomes depending on the certainty direction. If he is certain about choosing the correct aisle (high certainty of being correct and therefore confidence close to one), he will walk down this aisle to search for the product closely. On the other hand, if he is certain he made an incorrect decision (high certainty of being incorrect and, therefore, confidence close to zero), he will turn around and walk to another aisle. Another compelling example of the relationship between confidence and bidirectional certainty is the “change of mind” phenomenon that has been linked to confidence declining below 0.5 during the execution of the decision (Resulaj et al., 2009; van den Berg et al., 2016).

Although as exemplified, both certainty directions might result from the same decisional context, it was not until recently that they were studied concomitantly (Boldt & Yeung, 2015; Fleming and Daw, 2017; Yu et al., 2015). Prior to this, the assessment of information associated with erroneous decisions was extensively studied in the context of the *error monitoring and detection*, typically with binary reports (e.g. Charles et al., 2013; Rabbitt, 1966; Rabbitt & Rodgers, 1977; Yeung & Summerfield, 2012), whereas previous *confidence evaluation* studies considered mainly the graded certainty of being correct (e.g. Fleming et al., 2012; Hebart et al., 2014; Heereman et al., 2015; Kepecs and Mainen, 2012; Kiani and Shadlen, 2009). These studies provided important knowledge about the metacognition of perceptual, memory-based and value-based decision-making (e.g. Fleming and Dolan, 2012a; Hampton, 2001; Kiani and Shadlen, 2009; Monosov and Hikosaka, 2013).

Nonetheless, even the studies that addressed both certainty directions did not dissociate the relative influence of certainty in correct or incorrect decisions on confidence reports and metacognitive ability. Moreover, although research on different animal species implicated non-language-related cognitive processes in a computational framework of metacognition (Kepecs and Mainen, 2012; Kiani and Shadlen, 2009; Meyniel et al., 2015; Pouget et al., 2016), the ability to use certainty of being correct and certainty of being incorrect *implicitly*, without verbal confidence scale formulations, has not been measured in the same experiment.

The aim of this study was to capture probabilistic readouts that could provide separate quantifiable measures of each certainty direction without relying on explicit verbal formulations, and to test the hypothesis that implicit assessments of both certainty of being correct and certainty of being incorrect result in more adaptive post-decisional behaviors. To this end, we designed a novel experiment in which trials with post-decision wagering (PDW; Persaud et al., 2007) were interleaved with trials in which subjects were instructed to bet money *before* the perceptual decisions (Pre-decision wagering, PreDW). As briefly described here and in detail in the *Methods* (section 2.6), we compared PDW and PreDW in order to isolate the trial-specific decisional information used in readouts of certainty of being correct and certainty of being incorrect, and to quantify these readouts.

To tease apart these trial-specific readouts, we relied on the design in which in both PreDW and PDW trials subjects could use information about expected difficulty presented at the beginning of each trial to choose their wagers (Fig. 1). Moreover, subjects’ psychological biases were expected to have similar influence on their wagering behavior in both PreDW and PDW. For example, loss-averse subjects were expected to wager lower than non-loss-averse subjects.

**Figure 1.**
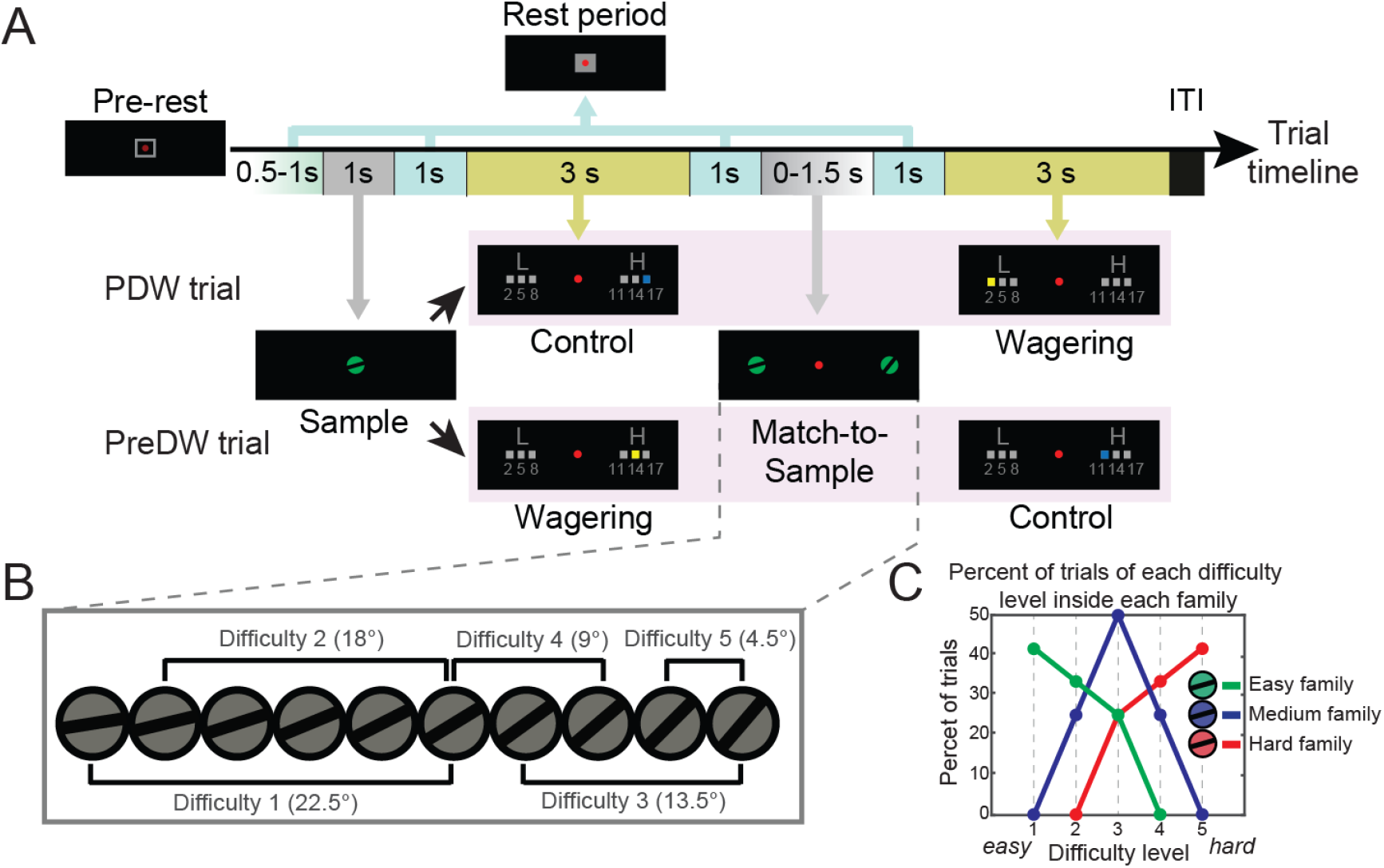
(A) Task design. The appearance of a red spot (for eye fixation) and of a gray framed-square (indicating buttons status) in the screen center signaled the trial start (pre-rest). The brightening of the red spot and the onset of the gray filled square indicated that subjects correctly adopted the rest position. The sample was then presented in the center of the screen for 1 s. During PDW trials, subjects first performed the control task. The letters H (‘high’) and L (‘low’) were presented on each side of the screen. The presentation sides varied randomly. A blue square appeared above a specific “wager” and subjects had first to select high or low and then use the same button repeatedly to select the instructed “wager” option. The selection always moved from center-out. Overall, subjects had 3 s to select the instructed “wager”. Then, subjects performed the match-to-sample task by choosing the image they believed was the match. Next, subjects performed the actual PDW wagering task, which was similar to the control task except that, after freely choosing high or low wager category, subjects could move the yellow square that appeared above one of the three specific wagers to select a desired option. PreDW trials were similar to PDW trials, with the difference that the wagering task and the control task order in the trial timeline was reversed. (B) Five difficulty levels were created by different orientation contrasts between the match and the non-match (linearly from 4.5° to 22.5°). (C) Proportion of trials in easy family (green), medium family (blue) and hard family (red) from each difficulty level (1 to 5).

On the other hand, only when subjects wagered after the perceptual decision (PDW trials) they could add trial performance-specific information - such as the certainty level and the certainty direction associated with the trial decision - to the non-trial-specific information in order to wager more adaptively. Hence, we subtracted certainty-related PreDW measures (based on non-trial-specific information) from certainty-related PDW measures (based on both trial-specific and non-trial-specific information) to isolate the PDW trial-specific information used during certainty readouts (Fig. 2). Thus, the use of PreDW baseline, which accounted for non-trial-specific biases and expectations, was instrumental for isolating the readouts of certainty during PWD. Using this approach, we calculated certainty of being correct by comparing PreDW and PDW measures associated with *correct* perceptual decisions. And, separately, we calculated certainty of being incorrect by comparing PreDW and PDW measures associated with *incorrect* perceptual decisions.

**Figure 2.**
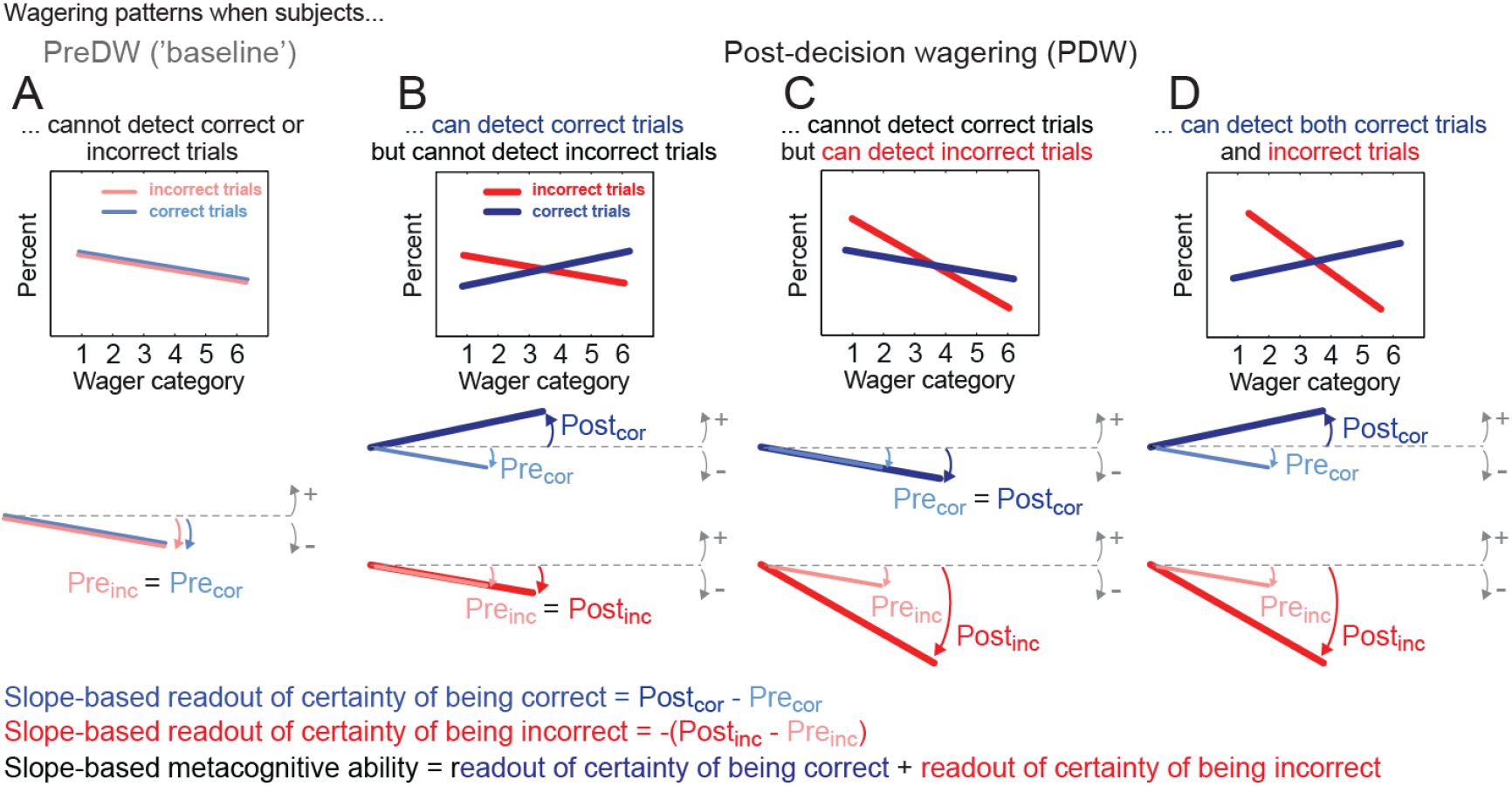
A proposed framework for calculating readouts of certainty of being correct and certainty of being incorrect based on the slopes of linear fits to wager-specific proportions of correct and incorrect trials. (A) Pre-decision wagering (PreDW) baseline. Slope-correct (the angle between the light blue linear fit and the horizontal plane) and slope-incorrect (angle between the light red linear fit and the horizontal plane) when subjects cannot detect correct or incorrect trials (no decision yet has been made). Since subjects cannot tell correct from incorrect trials, each wager has the same proportion of correct and incorrect trials and therefore the slopes should be the similar (*Pre*_*inc*_ *= Pre*_*cor*_). In this example, subjects use low wagers more often than high wagers (e.g. because they are risk-averse, or have a low overall confidence in their performance), which leads to negative baseline slopes. (B) Similarly, post-decision wagering (PDW) slope-correct (*Post*_*cor*_, blue) and slope-incorrect (*Post*_*inc*_, red) when subjects can only detect correct DMTS decisions. PDW slope-correct is larger than PreDW slope-correct, indicating the readout of certainty of being correct, which is measured by subtracting PreDW slope-correct (*Pre*_*cor*_) from PDW slope-correct (*Post*_*cor*_). The readout of certainty of being incorrect is still at zero level (*Pre*_*inc*_ *= Post*_*inc*_). (C) PDW slope-correct and PDW slope-incorrect when subjects can only detect incorrect trials. PDW slope-incorrect (*Post*_*inc*_) is smaller than PreDW slope-incorrect (*Pre*_*inc*_), indicating the readout of certainty of being incorrect. (D) PDW slope-correct and PreDW slope-incorrect when subjects can detect both correct and incorrect perceptual decisions. *Post*_*cor*_ and *Post*_*inc*_ are different from *Pre*_*cor*_ and *Pre*_*inc*_, respectively. In this case, slope-based metacognitive ability (the sum of both certainty readouts) is influenced both by the readout of certainty of being correct and by the readout of certainty of being incorrect.

We found that subjects indeed wagered accordingly with non-trial-specific information during PreDW trials, whereas they used trial-specific information to bet money more efficiently during PDW. Importantly, the trial-specific information was used to read out both certainty directions. To our knowledge, this is the first measurement of the bi-directional assessment of metacognitive information (readouts of certainty of being correct and certainty of being incorrect) based on implicit reports and using the same paradigm. Our measurements showed that humans are able to assess whether they have made correct or incorrect decisions and use this information to behave adaptively without the need of verbal formulations. Moreover, although the readouts of certainty of being correct influenced subjects’ earnings, it was the ability to read out certainty of being incorrect that affected most the amount of money earned during PDW. These results demonstrate the importance of readouts of certainty of being incorrect in confidence judgments and suggest their adaptive value for the future actions.

## 2. Methods

### 2.1 Subjects

Eighteen subjects (6 males; mean age 25.7 years) were recruited via an online platform of the University of Goettingen. All subjects had normal or corrected-to-normal vision. Data from one subject was discarded because of insufficient number of trials in some of the conditions. Subjects were paid according to their performance (please see below). The experimental procedures were approved by the local ethics committee.

### 2.2 Experimental setup

Subjects sat in front of an LED screen (1600 x 1200 resolution) at 51 cm viewing distance and responded manually using two capacitive proximity sensors (buttons) connected to the computer via parallel port. Subjects positioned their head over an adjustable chin rest and had their head fixed with an adjustable strap for better stabilization. Gaze position was acquired with 60 Hz miniature infrared eye tracker camera and ViewPoint 2.8.6.21 software (Arrington Research). The task was controlled via MATLAB (Mathworks Inc) using the Psychophysics toolbox (http://psychtoolbox.org/). Subjects performed practice trials until they became familiar with this experimental setup.

### 2.3 Delayed match-to-sample task

Subjects completed 360 trials of a visual delayed match-to-sample (DMTS) task in which they had to find, between two options, the match for a preceding sample, which consisted of one gray circle of 1.5° of visual angle radius with an oblique black bar crossing its center (Fig. 1). Ten different sample options were generated by varying the bar orientation in counterclockwise rotation from the horizontal plane (from 18° to 58.5°). One of these ten samples was presented pseudo-randomly at the beginning of each trial in the center of the screen. During match-to-sample presentation, one sample-like image was presented 9° to the right and another one 9° to the left of the center of the screen (eye fixation spot). One of them had a bar in the same orientation as the sample (match) and the other one had a bar in a different orientation (non-match). Subjects responded by using the button of the hand positioned in the same side of the screen of the image selected as the match (Fig. 1A). Five difficulty levels were created by different orientation contrasts between the match and the non-match (from 4.5° to 22.5°; Fig. 1B). Trials with different difficulty levels were grouped into three families. The overall level of difficulty of each family was determined by the different proportions of trials of each difficulty level. The sample color - green, blue or red – cued these families: easy, medium or hard, respectively (Fig. 1C).

Subjects were informed before the experiment that colors were related to different levels of difficulty, but they were not told about the link between specific colors and difficulty of each family.

### 2.4 Pre-decision wagering, post-decision wagering and the control task

The metacognitive decision was a wagering task in which subjects were asked to bet money on the correctness of their perceptual decision. They won the wagered money for correct DMTS decisions and lost it for incorrect DMTS decisions. In half of the 360 trials, subjects wagered after the DMTS decision (Post-decision wagering, PDW) and in the other half of the trials subjects wagered before the DMTS decision (Pre-decision wagering, PreDW). PreDW and PDW trials were pseudo-randomly interleaved. PreDW was used as baseline condition in further analyses (see section 2.6). During wagering, subjects made the metacognitive decisions by selecting first to wager high (wager categories 4, 5 and 6) or low (wager categories 1, 2 and 3), and afterwards by selecting a specific wager category among low or high options. Subjects had to select a specific wager within 3 s by using the button of the hand positioned in the same side of the corresponding selected option. This two-stage wager selection procedure was used so that we could assess reaction times (see *section 3.6*) despite the graded response scale used for metacognitive decisions: the first stage binary response was intended to preclude additional influences on the reaction times due to variability in the urgency to select a specific wagering option along the scale.

In addition to the wagering, we used a control task, in which subjects had to select a visually-cued response option. This task worked as an “instructed” wagering (Fig. 1) and did not influence subjects’ earnings. It aimed to equalize, across PDW and PreDW trials, the cognitive effort due to intervening distractions (visual stimulation, object selection, and corresponding time interval). On PDW trials, subjects performed the control task before the DMTS decision, at the same period they were wagering on PreDW trials, and vice-versa for PreDW trials.

Subjects started the experiment with 10 Euros and could earn up to 30 Euros according to their performance. They wagered on the correctness of every DMTS decision using the following pay-off matrix, which was explained to them before the experiment:

**Table 1.** Wagering payoff matrix.

| DMTS decision | Low wagers |  |  | High wagers |  |  |
| --- | --- | --- | --- | --- | --- | --- |
| Correct | 2 cents | 5 cents | 8 cents | 11 cents | 14 cents | 17 cents |
| Incorrect | -2 cents | -5 cents | -8 cents | -14 cents | -17 cents | -20 cents |

As can be seen from the pay-off matrix, if subjects wagered low, they were rewarded and punished in the same way for correct and incorrect perceptual decisions, respectively. But when they wagered high, their incorrect perceptual decisions were punished with 3 cents more than they would have earned for correct DMTS decisions. This pay-off matrix was designed during pilot experiments in which subjects reported that they knew they were performing generally above the chance level (50%) and thus could earn money by simply wagering high all the time. To counteract this strategy, we encouraged subjects to evaluate every perceptual decision by punishing high wagers associated to incorrect DMTS decisions more than low wagers.

### 2.5 Trial timeline

Eye and hand movements were controlled throughout the trial. Each trial started with the appearance of a red sport and a gray framed-square in the center of the screen. Subjects were positioned in the rest position when they fixated the gaze inside the eye fixation window (3° visual angle radius around the red spot) and, concomitantly, positioned the right and left thumbs over two separate buttons. After a variable delay in the rest position (0.5-1 s), the sample was presented in the center of the screen for 1 s. After sample presentation, subjects had to maintain the rest position for another 1 s before the control task (for PDW trials) or the wagering task (for PreDW trials). Another period of 1s separated control/PreDW from the match-to-sample task. Subjects had up to 1.5 s to select the image they believed was the match. After the perceptual decision report and another interval of 1 s, subjects performed the wagering task (PDW trials) or the control task (PreDW trials, Fig. 1A).

There was no trial-by-trial feedback about the correctness of match selection. One feedback about the overall earnings collected so far was presented during a break that occurred after 180 complete trials, and the final earned value was presented after 360 complete trials. Trials in which subjects broke eye or hand fixation requirements, or were too slow to respond in one of task response periods, were aborted and repeated at a later time.

### 2.6 Slope-based measurements for certainty of being correct and incorrect: importance of PreDW baseline

We developed a new approach (named *slope-based measurements*) to calculate separate readouts of certainty of being correct and certainty of being incorrect. These measurements are based on linear fits derived from the proportions of correct or incorrect perceptual decisions that each of the 6 wagers was assigned to. These proportions were calculated by dividing the number of correct trials each wager was assigned to by the total number of correct trials (wager-specific proportion of correct trials) and, separately, dividing the number of incorrect trials each wager was assigned to by the total number of incorrect trials (wager-specific proportion of incorrect trials). For example, if a subject assigned the lowest wager to 15 of the 30 incorrect trials, this wager’ s proportion of incorrect trials is 50%. If the second lowest wager was assigned to 9 of the 30 incorrect trials, its proportion of incorrect trials is 30%, so on and so forth. Next, we fitted a linear regression to the 6 wager-specific proportions of correct trials and another linear regression to the 6 wager-specific proportions of incorrect trials. The slopes of those fits were named “slope-correct” and “slope-incorrect”, respectively, and were associated with the ability to read out each certainty direction. Hence, subjects whose proportions of correct trials increased towards the highest wager would demonstrate, through this positive slope-correct, the ability to read out certainty of being correct (see Fig. 2B).

However, a problem of using this approach for PDW in isolation is that, in addition to trial-specific information, those wager-specific proportions might also be influenced by unspecific factors such as the general task difficulty and psychological biases (e.g. loss aversion or overconfidence). For example, subjects with higher loss aversion and/or facing harder perceptual difficulty might choose low wagers more often than high wagers, independently of their performance in the preceding decision. In this case, the resulting slope would not be zero, but negative, even without any assessment of trial-specific performance. In order to disentangle the influences of such unspecific factors from the assessment of trial-specific performance, we used the PreDW task as the baseline for the slope-based measurements. During PreDW, subjects could only access their average performance in each of the three difficulty families, which were indicated by the sample color (green, blue or red for the three families: easy, medium or hard, respectively, Fig. 1C). Importantly, subjects were not able to predict, at the moment they were wagering, their actual trial performance. Consequently, subjects should end up assigning the wagers randomly to correct and incorrect trials, generating similar PreDW slope-correct and PreDW slope-incorrect values. This similarity is a requirement for the baseline condition, and it is present in our results (see *section 3.2* for further information).

During PDW, on the other hand, subjects might have access to their trial-specific DMTS performance. The better they assess this information (i.e. metacognitive ability), the more PDW slope-correct and PDW slope-incorrect become distinct from each other and from baseline PreDW slopes. This happens when subjects are able to assign high wagers more often to correct trials and low wagers more often to incorrect trials, *relative* to the PreDW baseline. The difference between PDW slope-correct (*Post*_*cor*_) and PreDW slope-correct (*Pre*_*cor*_) characterizes the readout of certainty of being correct. When subjects are able to detect correct trials, *Post*_*cor*_ is larger than *Pre*_*cor*_ (Fig. 2B). The same calculation is done independently for incorrect DMTS decisions, with inverted assumption: when subjects are able to detect incorrect perceptual decisions, *Post*_*inc*_ is smaller than *Pre*_*inc*_ (Fig. 2C).

Subjects’ metacognitive ability (the ability in detecting correct and/or incorrect DMTS decisions on a trial-by-trial basis) will be reflected in the sum of their abilities to read out certainty of being correct and certainty of being incorrect. The metacognitive ability reaches highest levels when subjects are able to read out both certainty directions (Fig. 2D).

It is important to emphasize that, although we use words “identify” and “detect”, we believe that reading out certainty is a probabilistic process. Therefore the readouts reflect the detection of correct and incorrect perceptual decisions in a probabilistic manner (Pouget et al., 2016).

### 2.7 D-prime (d′) and meta-d′ calculation, and relationship to slope-based metacognitive ability

In the section 3.5 of Results we compare the slope-based metacognitive ability with an established measure of metacognitive ability, meta-d*’* (Maniscalco and Lau, 2012). Meta-d*’* was calculated using the parameters of the Signal Detection Theory model as applied in Maniscalco and Lau (2012) code available online (http://www.columbia.edu/∼bsm2105/type2sdt/archive/index.html). This method estimates the value of perceptual (Type 1) sensitivity (d′) that would have been required to produce the observed metacognitive hits and false alarms (Type 2 sensitivity). Since the wager-specific proportions of correct and incorrect trials, or in other words the accuracy-conditional probabilities of using each wager, are also used for the calculation of the standard Type 2 receiver operating characteristics, ROC, the same data go into the calculation of meta-d′ and into the calculation of the slope-based *PDW-only* estimates. Therefore, these two measures are closely related (in our data they present a correlation of R=0.9, p<0.0001; imperfect correlation might be due to meta-d′ being dependent on the Type 1 criterion, and because it uses a probit regression while the slope-based analysis uses linear regression).

Nevertheless, the main difference between the meta-d′ and the slope-based metacognitive ability is that the latter is predicated upon the comparison between PDW vs. PreDW and allows (as our results suggest) distinguishing between the two certainty directions. Meta-d′, on the other hand, is calculated using only PDW but cannot distinguish between the two certainty directions. The major advantage of meta-d′ is that it is calculated on the same scale as d′ and, therefore, these two measures can be directly compared. The metacognitive efficiency, defined as meta-d′ divided by d′ (meta-d′/d′), reflects the comparison between how well subjects used the information available for metacognitive decisions and for perceptual decisions (Maniscalco and Lau, 2012).

Based on the d′ and meta-d′ estimates, we created two groups of metacognitive efficiency by dividing the subjects into those who performed better on metacognitive PDW decisions compared to perceptual DMTS decisions (high metacognitive efficiency group, meta-d′/d′>1) and those who performed better in the DMTS task than during PDW (low metacognitive efficiency group, meta-d′/d′<1). Since values of meta-d′>d′ would not be predicted if the same evidence is used for the Type 1 and the following Type 2 decisions (Maniscalco and Lau, 2012), we compared these two groups in order to investigate if a higher metacognitive efficiency could be associated with access to additional information that was not available for the Type 1 decision (Yu et al., 2015; Fleming and Daw, 2017).

### 2.8 Statistical analysis

We performed one-way, two-way and mixed-effects ANOVAs, linear correlations or t-tests using MATLAB (Mathworks Inc), as specified in the *Results*. R (The R Foundation) was used to perform multiple regression and linear mixed-effects regression models (R package nlme; Pinheiro et al., 2007). The mixed-effects ANOVAs and the linear mixed-effects regression models allowed us to include all data (unbalanced design) and still utilize repeated measures when appropriate. When required, post-hoc tests were performed and corrected using Bonferroni correction.

## 3. Results

17 human subjects were asked to carry out a visual perceptual decision of varying difficulty (delayed match-to-sample, DMTS task) and a wagering task either before (pre-decision wagering, PreDW) or after (post-decision wagering, PDW) the perceptual decisions. Trial types (PreDW and PDW) and difficulty levels were randomly interleaved.

### 3.1 Subjects performed similarly in the DMTS task during PDW or PreDW trials, and wagered according to the information available in each trial type

We performed a two-way ANOVA for repeated measures to assess if perceptual performance varied between trial types (PreDW and PDW, factor 1) and among difficulty levels (factor 2). As expected, subjects performed better in the DMTS task on trials of lower difficulty (mean±SE for difficulty levels 1 to 5: 86.7±3.3 81.4±2.2 75.0±3.0 67.8±2.4 49.7±2.7%; F_4,64_=51.439, p<0.0001). There was no difference in average DMTS performance between PreDW and PDW trial types (F_1,16_=2.104, p=0.17) and no interaction effect (F_4,64_= 0.970, p=0.43), showing that subjects performed similarly in PreDW and PDW trial types across the five difficulty levels.

We next tested, with separate linear mixed-effects regression models for PreDW and PDW trials, if there were differences in wagering behavior among the five difficulty levels or among the three families. As described in *Methods*, families had different proportions of trials of each difficulty level (Fig. 1C), and were signaled to subjects by the color of the sample. As expected, during PreDW trials subjects wagered according to the families (p<0.001 for all pair-wise comparisons between families), and not according to the actual trial difficulty level, which was unknown to the subjects at the moment they were wagering (p>0.05 for all difficulty levels; Fig. 3A). In PDW trials, although there was a significant difference between easy and hard families (p<0.05), this difference was driven by the actual difficulty levels (p<0.05 for the comparisons between all difficulty levels, except between difficulty levels 1 and 2; Fig. 3B).

**Figure 3.**
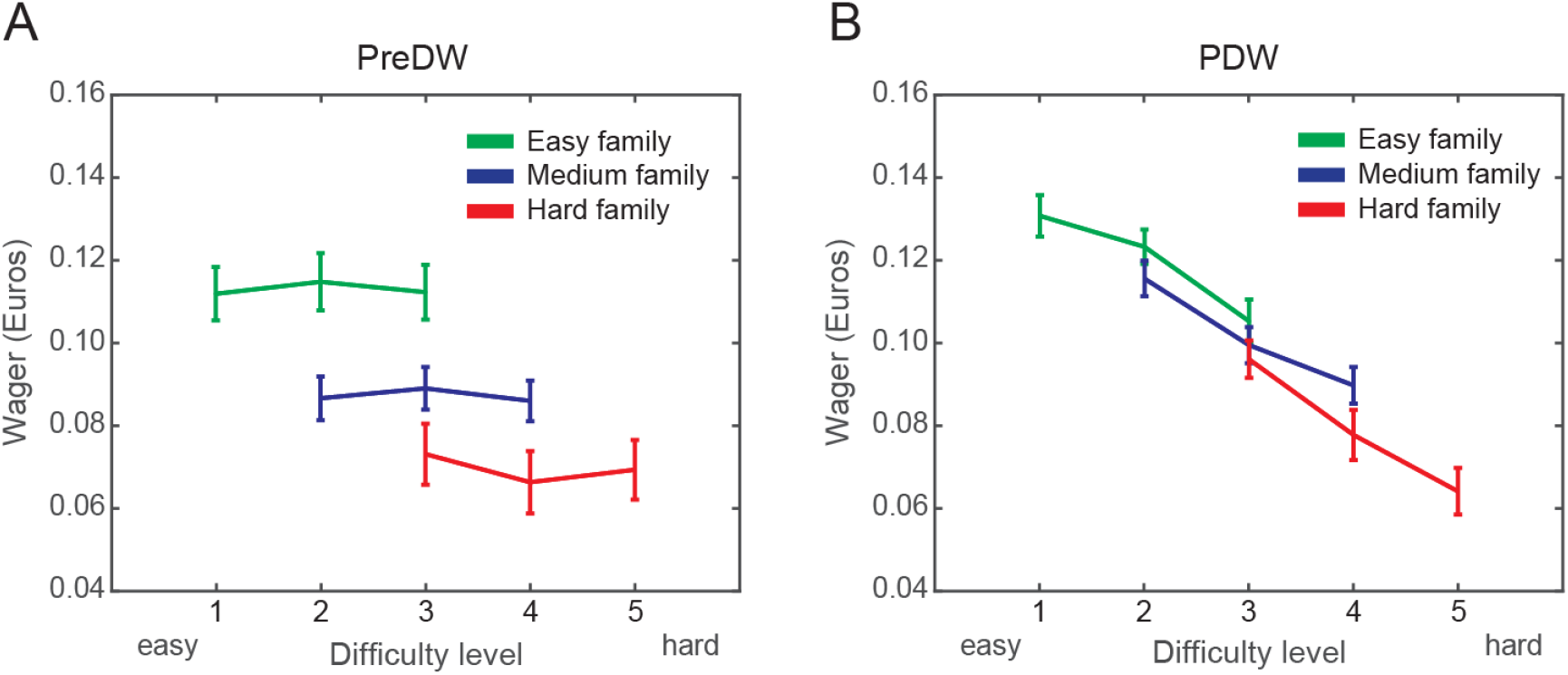
Means and standard errors of PreDW (A) or PDW (B) wagers for each perceptual difficulty level within each perceptual difficulty family: easy (green), medium (blue) and hard (red).

These results show that during PreDW subjects understood the differences among the families and wagered according to them. The results also indicate that during PDW trials subjects did not rely solely on the sample color (which signaled the average difficulty of each family). Instead, they also used trial-specific information accessed through the direct comparison between the two match options (trial-specific difficulty level).

With exception of d′ and meta-d′ calculations, the following results are based on measurements averaged across the five difficulty levels. In these measurements, we first averaged the results of different difficulty levels for one subject, and then we calculated averages across subjects. In the *section 3.7* we present data on separate difficulty levels to establish their relationship with subjects’ metacognitive readouts.

### 3.2 Slope-based measurements reveal that on average subjects read out both certainty directions

As described in the *section 2.6*, slope-correct and slope-incorrect are independent measures used to quantify readouts of certainty of being correct and certainty of being incorrect, respectively. These slopes are based on linear regressions fitted to the wager-specific proportions of correct and incorrect trials (Fig. 2). To isolate trial-specific readouts, we created a baseline condition derived from PreDW trials. PreDW provided us with general wagering trends that subjects might have developed based on non-trial-specific information (such as expected difficulty and psychological biases) which, when subtracted from PDW slopes, should provide slope differences resulting only from trial-specific information.

Since PreDW is based solely on non-trial-specific information, subjects should assign wagers randomly to the following correct and incorrect perceptual decisions and, in contrast to PDW, PreDW slope-correct should be similar to PreDW slope-incorrect (Fig. 4A). We used two-way ANOVA for repeated measures to test if slope-correct is different from slope-incorrect (factor 1) depending on the trial type (PDW or PreDW, factor 2). Independently of the other factor, PreDW slopes did not differ from PDW slopes (F_1,16_=0.007, p=0.93) but slope-correct differed from slope-incorrect (F_1,16_=64.039, p<0.0001). The interaction effect revealed that slope-correct and slope-incorrect were different depending on the trial type (F_1,16_=42.493, p<0.0001). Importantly, the post-hoc test showed that this difference occurred during PDW trials (t_16_=9.433, p<0.01; Fig. 4B), but not during PreDW trials (t_16_=2.204, p>0.05, Fig. 4A), suggesting that PreDW is a reliable baseline.

**Figure 4.**
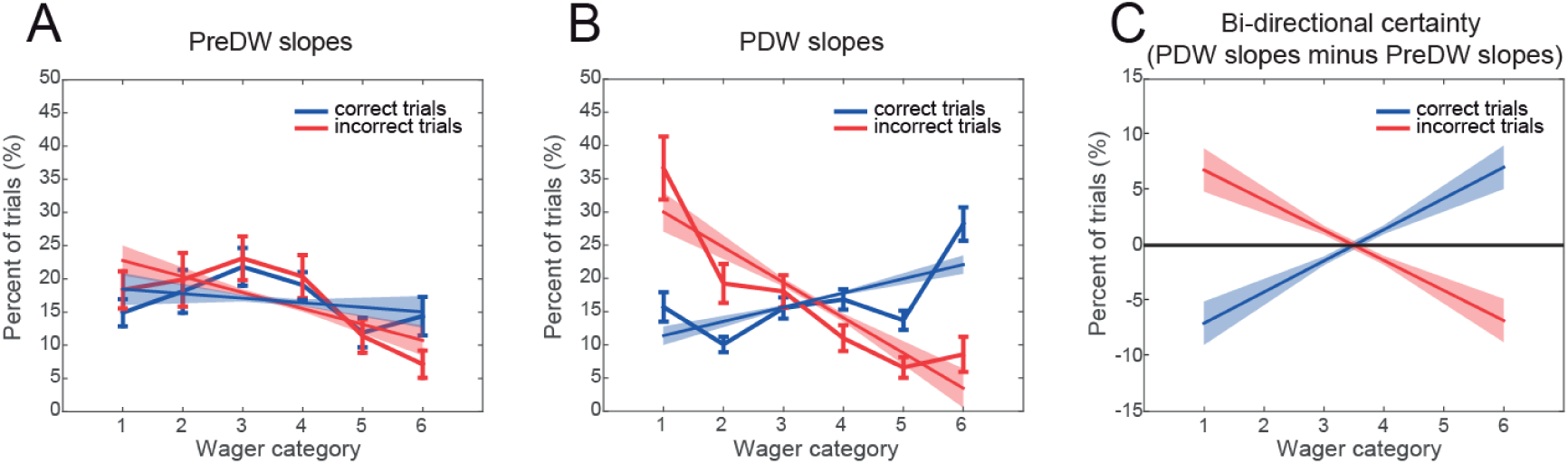
Means and standard errors of linear fits for correct trials (blue lines and shaded bands) and incorrect trials (red lines and shaded bands) for (A) PreDW (baseline) and (B) PDW, fitted to the data: means and standard errors of wager-specific proportion of correct trial (blue curves) and incorrect trials (red curves). (C) Mean and standard error of PDW slope-correct minus PreDW slope-correct (blue line and shaded band) and of PDW slope-incorrect minus PreDW slope-incorrect (red line and shaded band), for all subjects. See Supplementary Figure S1 for PreDW and PDW data plotted separately for each difficulty.

After verifying that we have a valid baseline, we tested if subjects were able to independently read out certainty of being correct and certainty of being incorrect, by calculating the difference between PreDW and PDW slopes separately for correct and incorrect decisions. PDW slope-correct was significantly higher than PreDW slope-correct (t_16_=3.6115, p<0.01), indicating that on average subjects read out certainty of being correct and used this information to wager more adaptively after the decisions. Similarly, PDW slope-incorrect was significantly more negative than PreDW slope-incorrect (t_16_=-3.7327, p<0.01). These slope-based measures indicated that on average subjects were able to use trial-specific information to avoid wagering high after incorrect perceptual decisions, while wagering high after correct decisions (Fig. 4C).

### 3.3 High and low metacognitive efficiency groups had similar performance in perceptual decisions but differed in metacognitive ability

In the present experiment, d′ (Type 1 sensitivity) reflects how well subjects identified the match during DMTS decisions; and meta-d′ (Type 2 sensitivity) reflects how well subjects used wagers to identify correct and incorrect DMTS decisions. We used Maniscalco and Lau (2012) method to measure d′ and meta-d′ on the same scale and to compare them directly (*section 2.7*). We plotted meta-d′ as a function of d′, and distinguished between two groups of subjects: a group of 11 subjects with meta-d′>d′, falling above the diagonal (high metacognitive efficiency group), and a group of 6 subjects with meta-d′<d′, falling below the diagonal (low metacognitive efficiency group; Fig. 5A).

**Figure 5.**
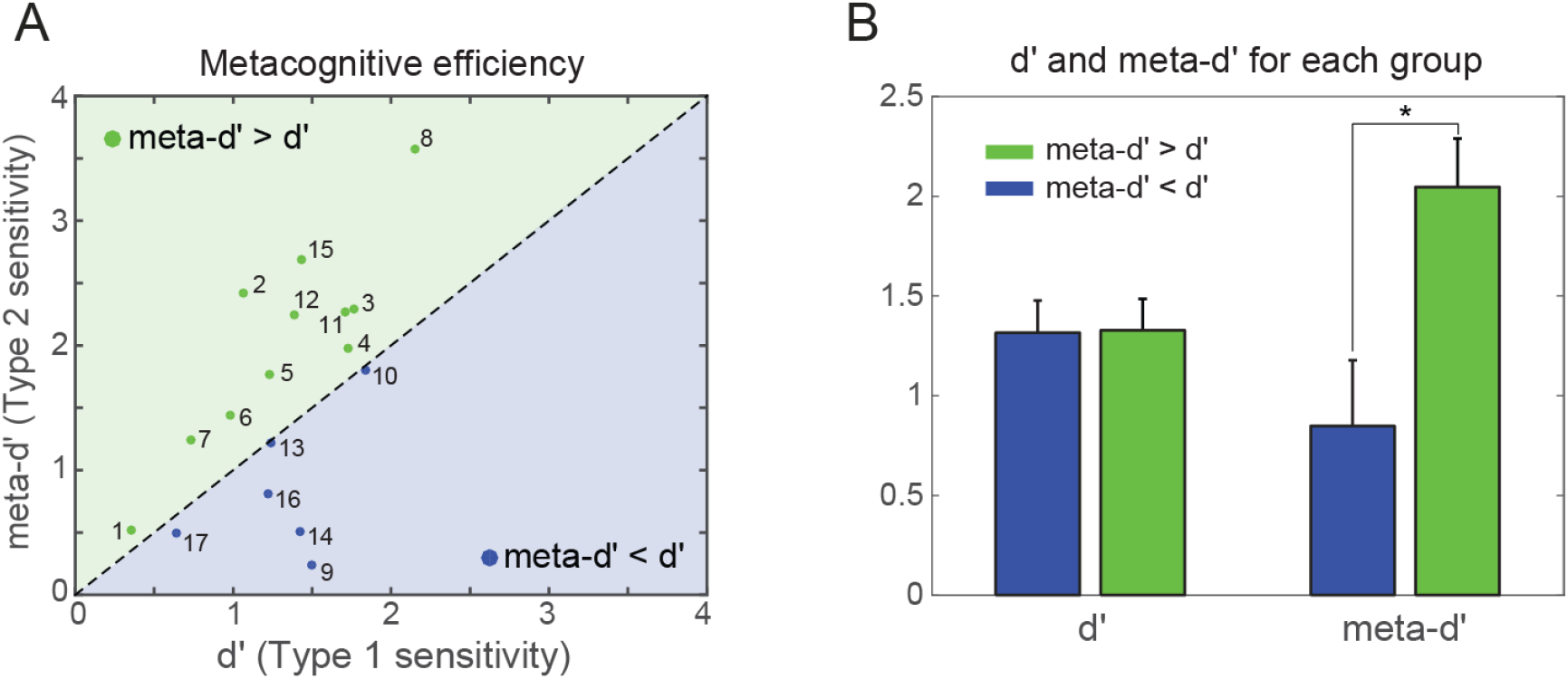
(A) Meta-d′ plotted as a function of d′. Eleven subjects with meta-d′>d′ (high metacognitive efficiency group) fell above the equality diagonal (green area), and 6 subjects with meta-d′<d′ (low metacognitive efficiency group) fell below the diagonal (blue area). (B) Means and standard errors of d′ and meta-d′ values for each group: meta-d′>d′ (green bars) and meta-d′<d′ (blue bars; *p<0.05).

Since we used a *post-hoc* grouping approach, it was important to check if the measurements (d′ and meta-d′) we used to create those groups varied significantly in the intergroup comparison. We applied a mixed-effect ANOVA with two factors: type of measurement (d′ and meta-d′, within-subjects) and group (meta-d′>d′ and meta-d′<d′, between-subjects). Meta-d′ was not different from d′ across the entire sample (F_1,15_=1.118, p=0.31), and there was no group difference averaging the two measurements (F_1,15_=4.256, p=0.06). However, the interaction effect was significant (F_1,15_=24.995, p<0.001). Post-hoc tests revealed that the two groups had the same d′ (t_15_=0.051, p=0.96), but the group of subjects with high metacognitive efficiency had higher meta-d′ than the group of subjects with low metacognitive efficiency (t_15_=3.198, p<0.05; Fig. 5B). This result allowed us to compare the two groups knowing that intergroup differences were not associated with differences in subjects’ performance in the DMTS task (Type 1 sensitivity), but only with their performance during PDW (Type 2 sensitivity).

### 3.4 Only the group of subjects with higher metacognitive efficiency read out certainty of being incorrect

To test if the differences between PDW and PreDW slopes which we found for all subjects together (*section 3.2*) were present in both low and high metacognitive efficiency groups, we performed two mixed-effect ANOVAs (PDW and PreDW slopes, within-subjects factor 1; groups, between-subjects factor 2), for slope-correct and slope-incorrect measures. The first ANOVA revealed that slope-correct was higher for PDW compared to PreDW trials (F_1,15_=10.218, p<0.01) without group difference (F_1,15_=0.017, p=0.90; meta-d′>d′: t_10_=2.791, p<0.02; meta-d′<d′: t_4_=2.892, p<0.05) or interaction between the factors (F_1,15_=0.433, p=0.52; Fig. 6A), indicating that group differences in metacognitive efficiency were not driven by the readouts about the certainty of being correct. The second ANOVA revealed that slope-incorrect was also different between PDW and PreDW (F_1,15_=11.070, p<0.01) without group difference (F_1,15_=1.825, p=0.20). However, the interaction effect was significant (F_1,15_=4.888, P<0.05) and the post-hoc test revealed that PDW slope-incorrect was different from PreDW slope-incorrect only for the high metacognitive efficiency group (t_10_=-4.742; p<0.01; Fig. 6A). Figures 6B and 6C further illustrate the difference between PWD and PreDW slopes for the two groups. Altogether, these results indicate that the difference between the groups in regard to the slope-based metacognitive ability, defined as the sum of readouts of certainties of being correct and incorrect (metacognitive ability 7±1 for the meta-d′>d′ group and 3±1 for the meta-d′>d′ group, non-paired t-test, t_15_=2.748, p<0.05), was due to the ability of subjects with high metacognitive efficiency to read out certainty of being incorrect.

**Figure 6.**
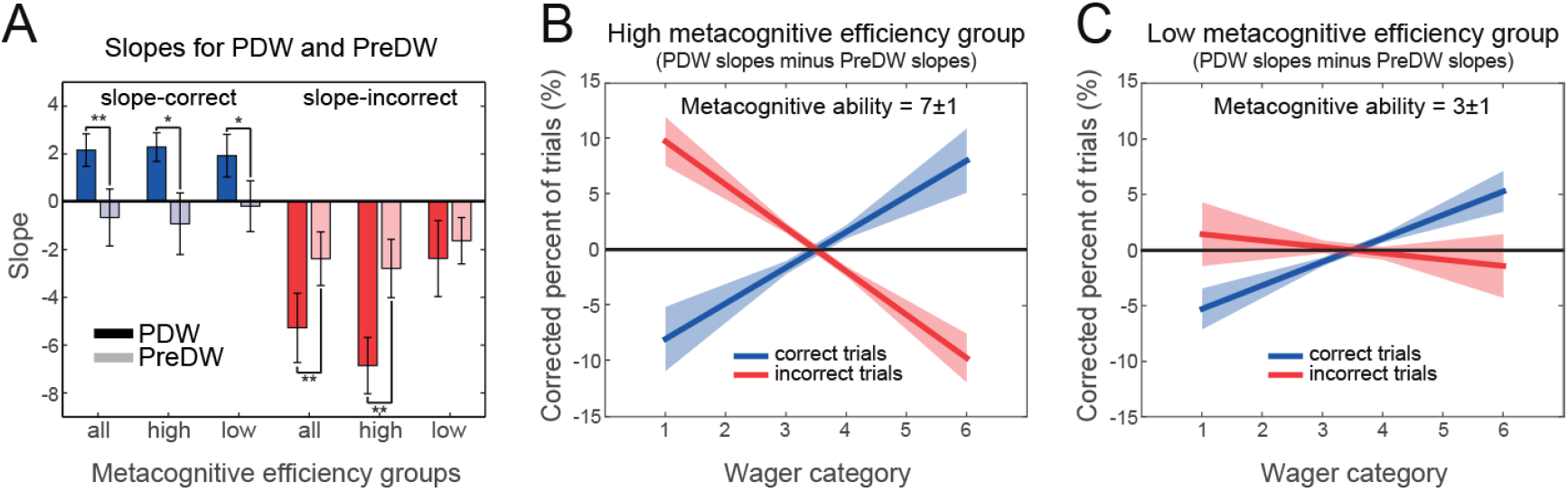
(A) Means and standard errors of PDW slope-correct (dark blue bars), PreDW slope-correct (light blue bars), PDW slope-incorrect (dark red bars), PreDW slope-incorrect (light red bars) from all subjects (“all”), high metacognitive efficiency group (“high”) and low metacognitive efficiency group (“low”; *p<0.05, **p<0.01 for differences between PDW and PreDW). (B) Mean and standard error of PDW slope-correct minus PreDW slope-correct (blue line and shaded band) and of PDW slope-incorrect minus PreDW slope-incorrect (red line and shaded band), for high metacognitive efficiency group. The text in the top of the panel shows mean and standard error of slope-based metacognitive ability. (C) Same as B, but for the low metacognitive efficiency group. All measurements represent data across difficulty levels (averaged within each subject) and then averaged across subjects.

The contribution of reading out certainty of being incorrect to metacognitive ability was further supported by the fact that several subjects from the high metacognitive efficiency group had perceptual performance *below the chance level* (50%) in the lowest wager category (Fig. 7A). To demonstrate this, we used mixed-effects ANOVA to test if the perceptual performance varied across the wager categories (within-subjects factor 1) and if this variation was similar between the high and low metacognitive efficiency groups (between-subjects factor 2). As expected, subjects performed better on trials in which they selected higher wagers (F_4,60_=23.837, p<0.0001). Importantly, the interaction effect showed that while the groups had the same general perceptual performance (F_1,15_=0.107, p=0.75), wager-specific perceptual performances were different between them (F_4,60_=7.077, p<0.0001; Fig. 7A). While the perceptual performance of the meta-d′<d′ group varied from 58.7% in the lowest wager to 82.1% in the highest wager, the meta-d′>d′ group had a range varying from *below chance* performance in the lowest wager (43.1%) to 94.7% in the highest wager. The perceptual performance in the wager category 1 for the meta-d′>d′ group was significantly below the chance level (t_10_=-2.569, p<0.05).

**Figure 7.**
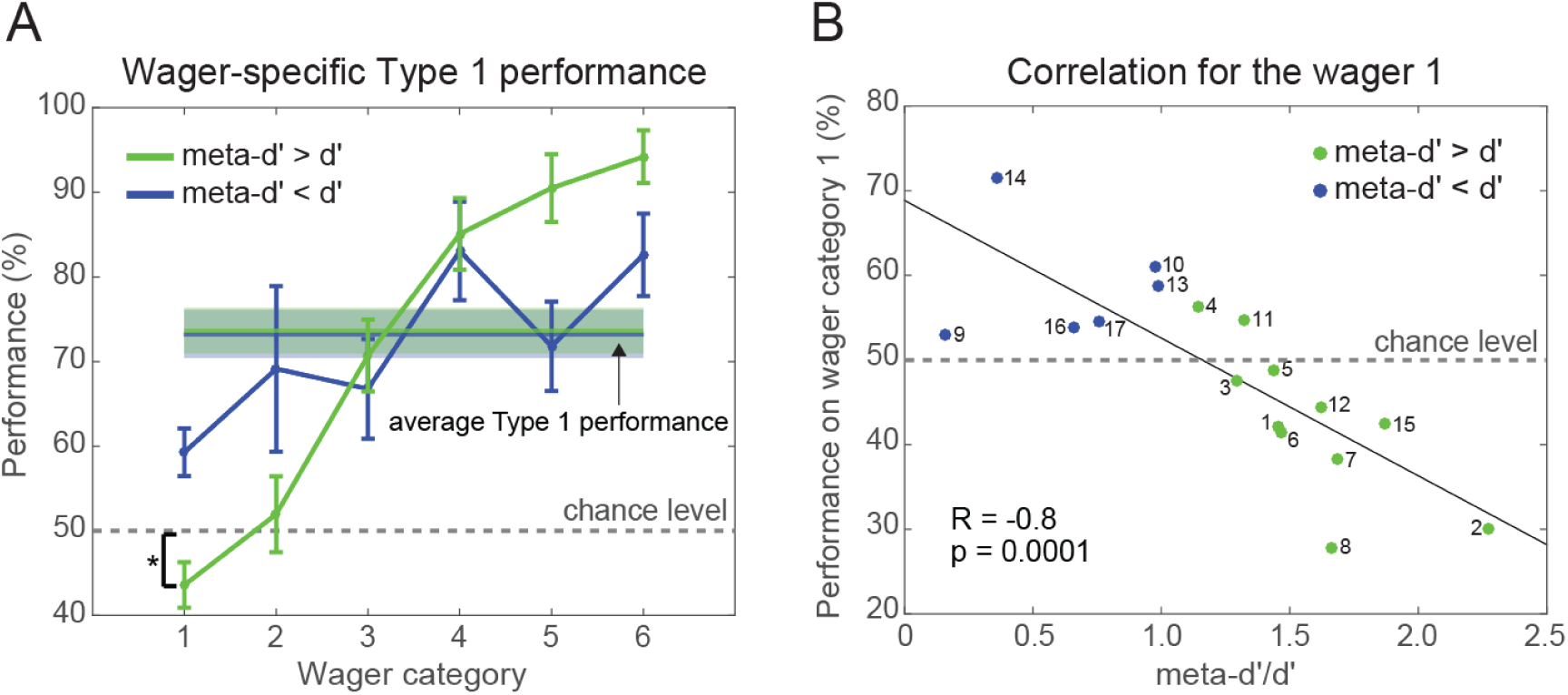
(A) Means and standard errors of perceptual performance for each metacognitive efficiency group (meta-d′>d′: green line and band; meta-d′<d′: blue line and band). Perceptual performance for the meta-d′>d′ group (green curve) is below chance level in the lowest wager (*p<0.05). The horizontal dashed black line represents the chance level (50%) for the DMTS task. (B) Scatter plot and the correlation between perceptual performance in the lowest wager category 1 and metacognitive efficiency (meta-d′/d′) across subjects (subjects from the group meta-d′>d′ are in green and subjects from the group meta-d′<d′ are in blue). The horizontal dashed black line represents the chance level for the DMTS task.

To further investigate this result, we calculated the correlation between perceptual performance in the wager category 1 and metacognitive efficiency across subjects (Fig. 7B). We found a strong negative correlation (R=-0.8, p<0.0001) between these two factors. This result is not surprising since meta-d′ is calculated based on accuracy-conditional probabilities of using each wager level (see *section 2.7*). However meta-d′ is influenced by *all* wager categories and it is noteworthy that the performance for wager category 1 is most correlated with meta-d′/d′ compared to performance in the other categories (second highest correlation was R=-0.56, data not shown), indicating a specific relevance of certainty about being incorrect for metacognitive efficiency. Most importantly, while several subjects with high metacognitive efficiency presented perceptual performance below the chance level for this wager category, none of the subjects with low metacognitive efficiency showed it. We interpret this result as a compelling demonstration that the meta-d′>d′ subjects were able to read out certainty of being incorrect to detect incorrect perceptual decisions and to assign the lowest wager to them.

### 3.5 Slope-based metacognitive ability is compatible with meta-d′

Since the slope-based measure that relies on the comparison of PreDW and PDW is an alternative approach we developed to assess separate contributions of certainty in being incorrect or correct to metacognitive ability, we compared it to meta-d′, an established and widely used measure. It is important to emphasize that the meta-d′ and the PDW-only slope-based calculations are based on the same data (see *section 2.7*), therefore we expect a good match between the slope-based and meta-d′ approaches, *provided* that the other component of the slope-based metacognitive ability, PreDW slopes, is indeed serving as a reliable baseline.

A strong positive correlation between the slope-based metacognitive ability and meta-d′ (R=0.88, p<0.0001) indicates that the slope-based approach can be considered a valid measure of metacognitive ability, with the advantage of allowing independent quantification of certainty of being correct and certainty of being incorrect readouts (Fig. 8).

**Figure 8.**
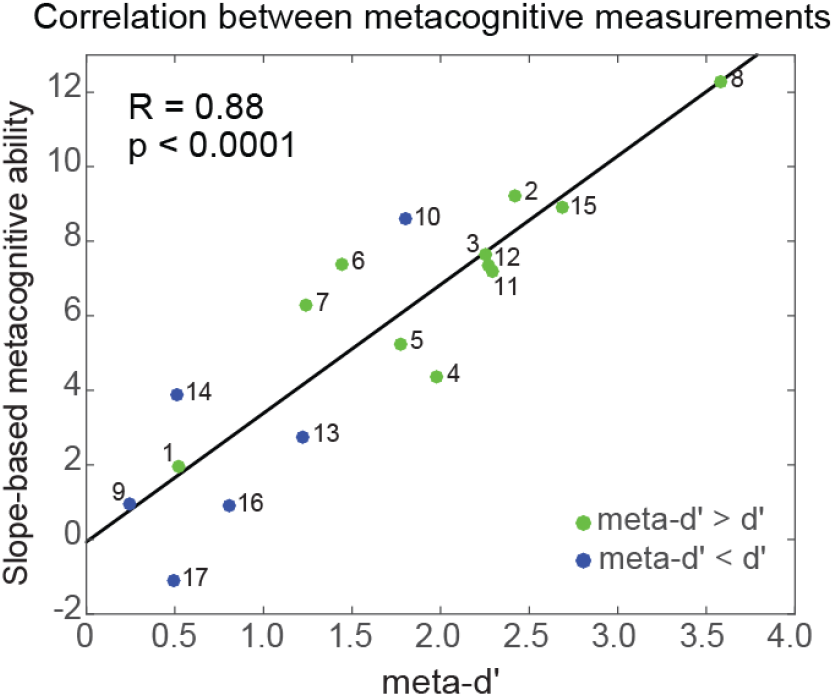
Correlation between two different measures of metacognitive ability. For each subject, the novel slope-based metacognitive ability measure is plotted against meta-d′ value. The black line is the best linear fit between the two measurements. Subjects are colored according to the metacognitive efficiency group (green, high metacognitive efficiency, blue, low metacognitive efficiency), numbers indicate subject labels.

The separate correlation between the slope-based readout of being correct and the meta-d′ was not significant (R=0.42, p=0.089), while the correlation between the slope-based readout of being incorrect and the meta-d′ was significant (R=0.57, p=0.016). In line with these findings, a multiple linear regression between meta-d′ and separate readouts of certainty in both directions showed a stronger influence of certainty of being incorrect on the meta-d′ (β =0.260, t_14_=6.495, p<0.001), as compared to certainty of being correct (β =0.211, t_14_=5.657, p<0.001).

### 3.6 Wager-specific metacognitive reaction times indicate increasing certainty in both directions of the wager scale

In the present experiment, each trial had two manual response periods. Perceptual reaction times (‘RT1’) reflect the time subjects took to report the DMTS decision. On average, subjects had faster RT1 when reporting easier DMTS decisions (F_4,60_=11.553, p<0.0001). Metacognitive reaction times (‘RT2’) reflect the time subjects took to choose between high or low wagers. Perceptual and metacognitive reaction times were also calculated separately for each wager (wager-specific RT1 and wager-specific RT2) to test the influence of perceptual reaction times on subsequent wagering behavior, as well as the relationship between certainty and metacognitive reaction times.

The mixed-effect ANOVA on perceptual RT1s (within-subjects factor 1: wagers, between-subjects factor 2: metacognitive efficiency group) showed that RT1s were associated with the subsequent selection of the wager (F_5,75_=21.140, p<0.0001). The association was unidirectional, with RT1s decreasing when followed by high wagers. The interaction effect showed that subjects from the high metacognitive efficiency group responded faster during perceptual decisions preceding some of the wagers (F_5,75_=4.004, p<0.01; Fig. 9A). There was no main effect of the group, perhaps because the low efficiency group included only 6 subjects (F_1,15_= 3.223, p=0.09).

**Figure 9.**
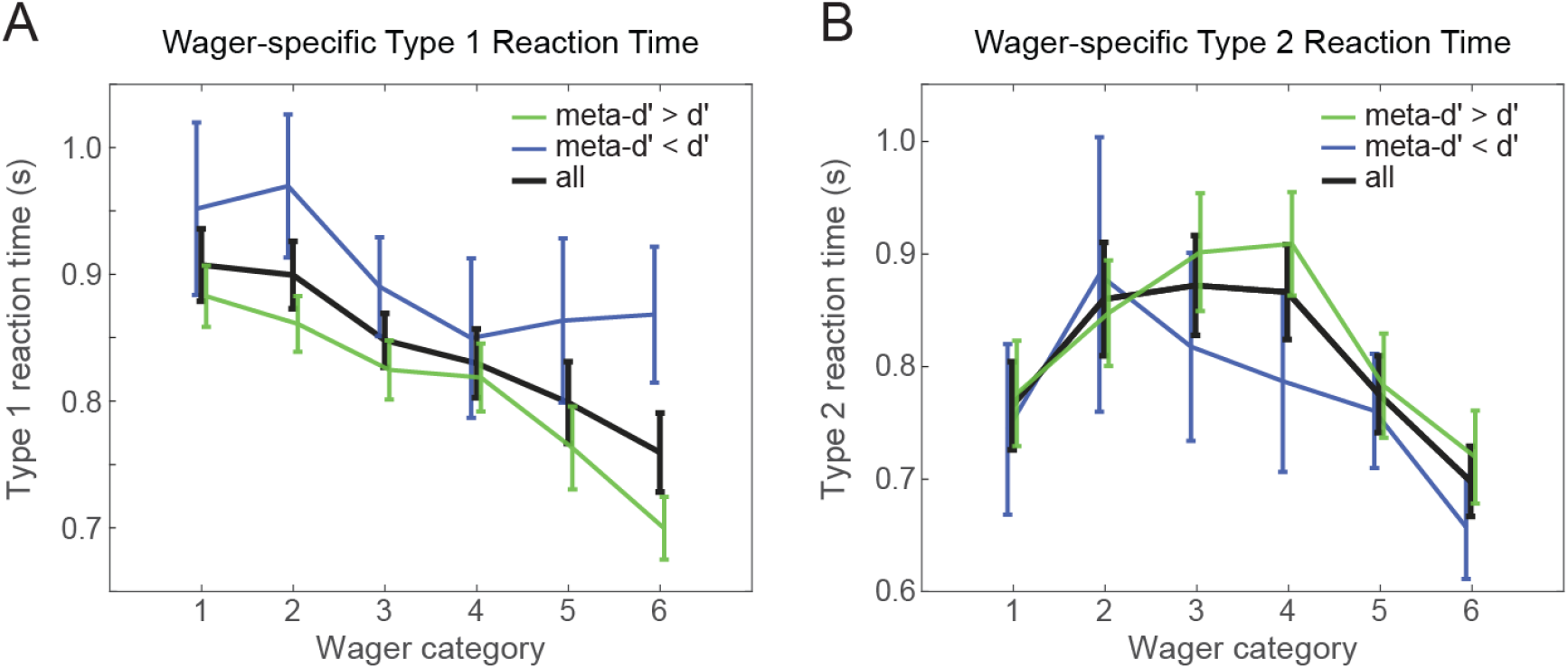
(A) Means and standard errors of wager-specific perceptual reaction times (RT1) across all subjects (black curve), for the high metacognitive efficiency group (meta-d′>d′, green curve) and for the low metacognitive efficiency group (meta-d′<d′, blue curve). (B) Means and standard errors of wager-specific metacognitive reaction times (RT2) across all subjects (black curve), for the high metacognitive efficiency group (green curve) and for the low metacognitive efficiency group (blue curve). All measurements represent averages across difficulty levels and across subjects.

The mixed-effect ANOVA on metacognitive RT2s (within-subject factor 1: wagers, between-subjects factor 2: metacognitive efficiency group) showed that mean wager-specific RT2s also differed among wagers (F_5,75_=10.087, p<0.0001). Across all subjects, RT2s were shorter in the wager categories 1 and 6 than in the middle wagers 3 and 4 (pair 1 and 3: t_16_=-3.514, p<0.05; pair 4 and 6: t_16_=2.546, p<0.05; Bonferroni-corrected), generating an inverted U-shape function of wager-specific RT2 (Fig. 9B). Since faster reaction times are associated with increased certainty (Kiani et al., 2014), this result further supports our hypothesis that certainty increases in both directions of the wager scale. We associate fast metacognitive reaction times in the wager category 1 with increased certainty of being incorrect, and fast metacognitive reaction times in the wager category 6 with increased certainty of being correct. But although the inverted U-shape seemed more pronounced in the high metacognitive efficiency group, the group comparison with mixed-effect ANOVA showed no significant differences between the groups (F_1,15_=0.397, p=0.54) or interaction between the factors (F_5,75_= 1.487, p=0.20; Fig. 9B). The group-specific pairwise comparison of wagers 1 and 3, however, revealed a significant RT2 difference only in the high metacognitive efficiency group (meta-d′>d′ group: t_10_=-4.495, p<0.01; meta-d′<d′ group: t_5_=-1.000, p>0.05) while the difference between RT2 for wagers 6 and 4 was significant in both groups (meta-d′>d′ group: t_10=_7.625, p<0.001; meta-d′<d′ group: t_5=_2.685, p<0.05). This result supports our interpretation of a link between bi-directional certainty and metacognitive reaction times.

### 3.7 Readouts of certainty of being correct and certainty of being incorrect increased with perceptual signal strength

For the analyses in preceding sections, we combined all five difficulty levels. Here, we performed two independent one-way ANOVAs for repeated measures to test for differences in certainty readouts across the difficulty levels. The first ANOVA showed that readouts of certainty of being correct decreased with difficulty level (F_4,64_=11.715, p<0.0001). The pair-wise post-hoc test revealed that these readouts decreased significantly only at the highest difficulty level (p<0.05; Fig. 10). The second ANOVA showed that readouts of certainty of being incorrect also decreased with increased trial difficulty (F_4,64_=4.529, p<0.01). The post-hoc test revealed that these readouts decreased significantly at the difficulty levels 4 and 5 (p<0.05, Fig. 10). These results show that subjects read out both certainty directions better when trials were easier. Considering that the probability of being incorrect increases in harder trials – and, therefore, in the opposite direction of certainty of being incorrect readouts – these results also suggest dissociation between the assessment of the DMTS difficulty and the trial-specific metacognitive assessment of the performance in the DMTS decision.

**Figure 10.**
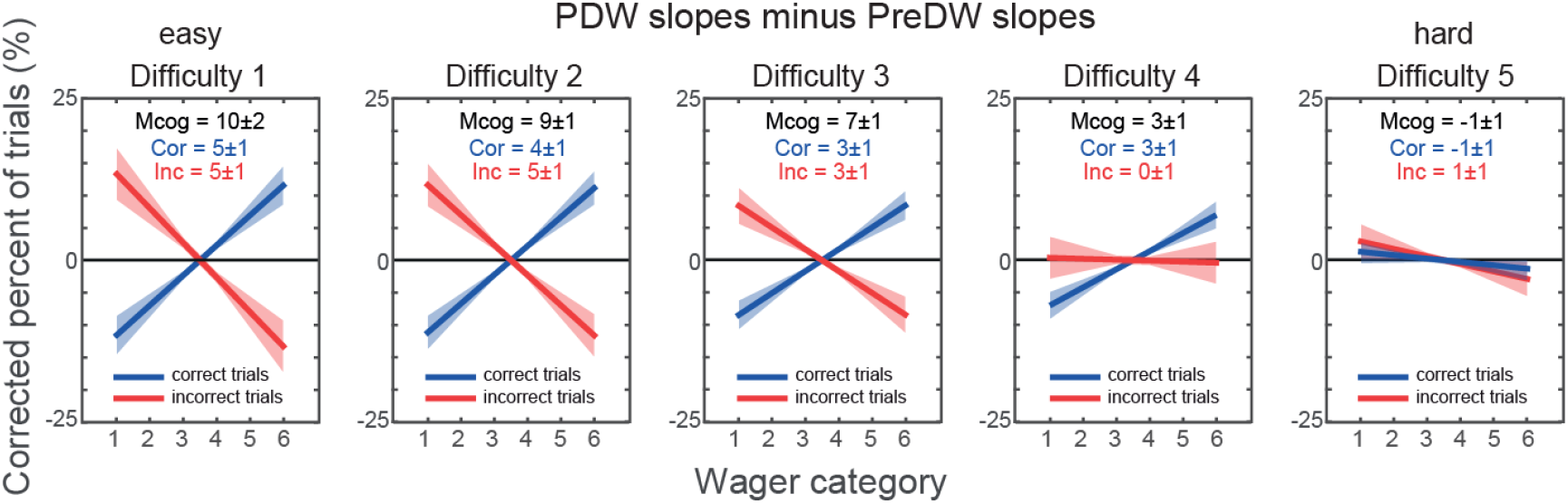
Means and standard errors of PDW slope-correct minus PreDW slope-correct (blue lines and shaded bands) and of PDW slope-incorrect minus PreDW slope-incorrect (red lines and shaded bands) for each difficulty level (easy to hard, 1 to 5). The text in the top of each panel shows means and standard errors of slope-based metacognitive ability (‘Mcog’), readout of certainty of being correct (‘Cor’) and readout of certainty of being incorrect (‘Inc’). See Supplementary Figure S1 for PreDW and PDW data plotted separately for each difficulty, used to derive those slopes.

### 3.8 Subjects with higher metacognitive ability earned more money in PDW, especially if they read out certainty of being incorrect

Finally, we used two linear regressions to understand how subjects’ PDW earnings were determined by their metacognitive abilities. The first linear regression showed that subjects’ earnings can be explained by their general metacognitive ability quantified by slope-based measurements (β =0.817, t_15_=3.686, p<0.005). A second multiple linear regression separated the two components of the slope-based metacognitive ability and showed that, although the ability to read out certainty of being correct partially explained subjects’ earnings (β =0.622, t_14_=2.646, p<0.05), their readouts of certainty of being incorrect influenced more how much they earned during PDW (β =1.060, t_14_=4.201, p<0.001).

## 4. Discussion

### 4.1 Bi-directional certainty readouts

In the present study we were able to measure trial-specific readouts of certainty of being correct and certainty of being incorrect by asking subjects to bet money (post-decision wagering, PDW) on their perceptual decisions (delayed match-to-sample task, DMTS). We quantified both certainty directions with a help of pre-decision wagering (PreDW), a task in which subjects could wager according to the average expected difficulty of the upcoming perceptual decision and their internal biases. The comparison between PDW and PreDW was utilized to isolate the trial-specific influences on the certainty readouts. On average, subjects were able to read out certainty of being correct and certainty of being incorrect, and to use this metacognitive information to bet money more efficiently.

These results also provided the first demonstration that the interpretation of implicit certainty scales should take into account certainty bi-directionality, since subjects utilized PDW to report bi-directional certainty readouts without explicit instructions to do so. PDW has been criticized for being highly influenced by individual biases associated with gains and losses (e.g. loss aversion and predisposition toward risky behaviors; Fleming and Dolan, 2010), but using a PreDW baseline allowed us to account for such biases and, consequently, overcome one of the main disadvantages of using wagering to access confidence.

In addition, we established a relationship between two directions of certainty readouts and subjects’ metacognitive efficiency (meta-d′/d′). Subjects with high metacognitive efficiency (i.e. those who performed better on metacognitive decisions compared to perceptual decisions, meta-d′>d′) and low metacognitive efficiency (meta-d′<d′) had on average the same performance on perceptual decisions, but only the high metacognitive efficiency group was able to read out both certainty of being correct and certainty of being incorrect, while the group of subjects with low metacognitive efficiency was able to read out only certainty of being correct. We argue in the next section that this group difference might be associated with additional information used during metacognitive decisions, which were not available at the moment subjects performed the perceptual decision. Nevertheless, considering that certainty of being incorrect influenced most how much subjects earned during PDW, these results suggest the adaptive value of readouts of certainty of being incorrect for the planning of post-decisional actions in situations in which immediate feedback is not available.

The certainty bi-directionality was further supported by the inverted U-shape function of the reaction times during metacognitive Type 2 decisions (RT2). In previous studies that used multiple-grade scales, Type 2 reaction times might not have been directly associated to certainty because the starting position of a cursor used for confidence reports was intentionally varied across trials to discourage advance motor preparation (e.g. Fleming et al., 2012b; Lebreton et al., 2015). Consequently, reaction times might have also depended on the starting position that could be closer or farther from the intended option, causing varying motor urgency not related to certainty. Conversely, in our task design, RT2s were based on initial binary decisions between two options randomly placed on each side of the screen (‘wager high or low’), and associated with specific wagers only at a second response step. This novel feature with two-step metacognitive decision reports allowed measuring RT2 association to certainty. Since faster reaction times are associated with increased certainty (Kiani et al., 2014), the inverted U-shape function of wager-specific RT2 suggests increased certainty in both directions of the wager scale.

### 4.2 Sources of additional information for readouts of certainty of being incorrect

The results of the slope-based approach demonstrated that subjects with high metacognitive efficiency were able to read out certainty of being incorrect (PDW slope-incorrect smaller than PreDW slope-incorrect). More than that, we interpret the fact that these subjects assigned the lowest wager more frequently to incorrect perceptual decisions than to the correct ones (cf. Figure 7) as a clear demonstration of their ability to recognize that they chose the wrong option in the DMTS task and use this information to avoid high losses. We speculate that such identification of incorrect choices might be associated with additional sources of information, resulting in improvement of the metacognitive performance in comparison to the preceding perceptual performance (meta-d′>d′).

When Maniscalco and Lau (2012) developed the calculation of meta-d′ at the same scale as d′, they initially assumed that the Type 2 sensitivity (meta-d′) should not exceed the Type 1 sensitivity (d′) because subjects use the same evidence in both types of decision. This assumption has been contested by more recent studies that show continuing post-decisional evidence accumulation when Type 2 decisions follow Type 1 decisions in time (Murphy et al., 2015; Yu et al., 2015, see also Fleming and Daw, 2017, for review). Along these lines, it can be argued that trial-specific readouts of certainty of being incorrect rely on additional information that was not available at the moment subjects committed to Type 1 decision, and that led to a reversal of the evidence accumulation direction towards the non-selected option (Yeung and Summerfield, 2012). Hence, metacognitive efficiency can be viewed not only as a measure of deleterious evidence leakage when meta-d′<d′, but also as a measure of extra evidence accumulation after Type 1 decisions, when meta-d′>d′.

Another non-mutually exclusive possibility is that certainty of being incorrect is a result of parallel metacognitive processing that runs concurrently with the formation of Type 1 decisions and thus, not necessarily predicated upon the post-decisional evidence accumulation. In this case, the dissociation between Type 1 performance and metacognitive readouts would be due to at least partially non-overlapping access to information available for the Type 1 and Type 2 decisions (Fleming and Daw, 2017). This view is supported by the presence of EEG “error-related negativity” before Type 1 responses (Gehring et al., 1993), and the simultaneous presence of decodable information about the required response and the actual (incorrect) response that subjects are preparing (Charles et al., 2014). Nevertheless, also in this account, certainty of being incorrect should be tightly linked to metacognitive sensitivity that exceeds Type 1 perceptual sensitivity (Fleming & Daw, 2017), which is in line with our results.

### 4.3 Evidence accumulation, trial difficulty and certainty scales

Here we bring together our results about certainty bi-directionality and trial difficulty, and the knowledge provided by previous work on confidence rating and published models of evidence accumulation (e.g. Pleskac and Busemeyer, 2010; Yeung and Summerfield, 2012; Yu et al., 2015). Note that while we only argue here within the post-decisional accumulation framework, some of the reasoning is also applicable to a parallel metacognitive processing that is dissociated from the perceptual evidence accumulation.

Figure 11 illustrates one hard (A) and two easy (B, C) *incorrect* trials of the DMTS task followed by six-grade certainty scale rating. The “evidence axis” exemplifies a theoretical range of evidence available for the task. On hard trials, the evidence is distributed narrowly around the Type 1 criterion. The easier the trial, the higher the probability that the evidence is accumulated further from the Type 1 criterion, and closer to the limits of the evidence axis. In a specific trial, if the post-decisional evidence remains in the same side of the Type 1 criterion as during the Type 1 decision, it is read out as certainty of being correct (blue curve). If the evidence crosses the Type 1 criterion, it leads to different levels of certainty of being incorrect (red curve). Hence, certainty increases towards the two directions of the evidence axis, resulting in a bi-directional certainty scale as the exemplified U-shape function of certainty at the bottom of the figure.

**Figure 11.**
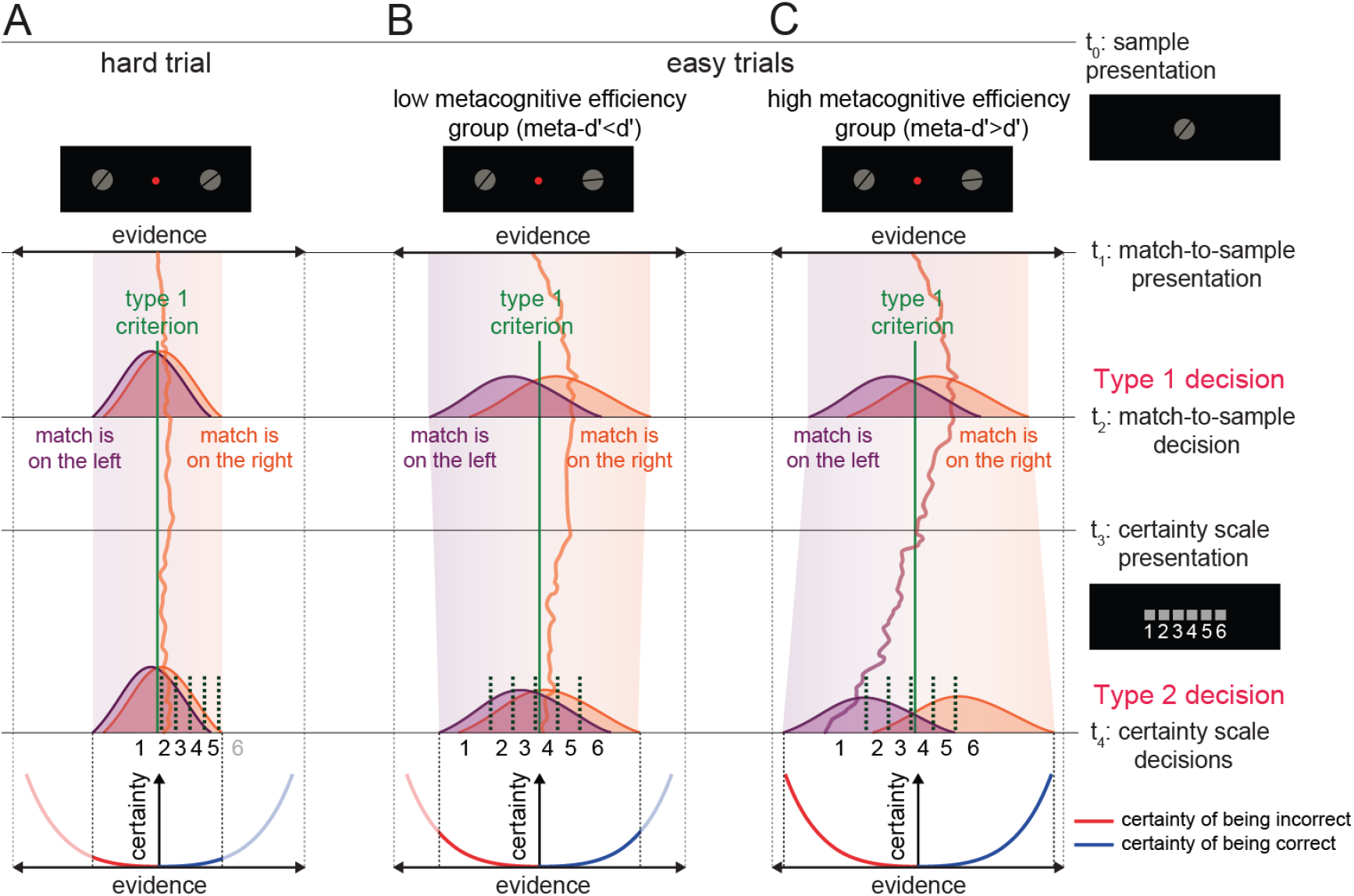
Certainty readouts in a post-decisional evidence accumulation framework. Trial time goes down from t_0_ to t_4_. Right (incorrect) option was selected during Type 1 decision. (A) On hard trials, d′ is small and the drift rate of evidence accumulation before and after the Type 1 decision is low, resulting in Type 1 performance at the chance level and low metacognitive ability. The Type 2 criteria distribution is closer to the Type 1 criterion, signifying low certainty. (B, C) On easy trials, the accumulation of evidence before the Type 1 decision is high and, consequently, d′ is large. One possibility is that the inter-subject variability in metacognitive efficiency arises from their ability to accumulate and read out post-decisional evidence. (B) Although subjects with low metacognitive efficiency have the same d′ as subjects with high metacognitive efficiency, they did not accumulate post-decisional evidence fast enough or failed in reading it out efficiently since their meta-d′ is smaller than d′. (C) Conversely, the post-decisional evidence accumulation is high for subjects with high metacognitive efficiency, allowing them to better distinguish correct from incorrect Type 1 decisions and increase their meta-d′. Compared to the low metacognitive efficiency group, the post-decisional accumulation of evidence is especially high when it drifts toward the non-selected match option, more strongly influencing the detection of the incorrect choices. Importantly, in this case, low ratings signify a high certainty (of being incorrect), rather than low certainty. Moreover, it is important to emphasize that in this illustration, Type 2 criteria distributions and the certainty directions would be flipped horizontally over the evidence axis in the case of Type 1 selection of the left option.

On hard trials (Fig. 11A), the difference between the perceptual evidence supporting each option is small, and consequently is d′. In our results, the Type 1 performance at the hardest difficulty level was at the chance level. Moreover, subjects did not show any metacognitive ability during Type 2 decisions in most difficult trials (Fig. 10). We therefore suggest that the post-decisional evidence accumulation rate remained as low as before the Type 1 decisions, and did not allow subjects to improve their metacognitive efficiency. On easy trials (Fig. 11B,C), the large difference between the perceptual evidence supporting each option of the Type 1 decision allowed subjects to better distinguish the match, yielding a high d′. Nevertheless, there were still some trials in which subjects selected the wrong option, as illustrated here. Our group comparisons indicate that subjects with low metacognitive efficiency did not accumulate post-decisional evidence or failed in reading it out efficiently, since their meta-d′ was smaller than d′ (Fig. 11B). We suggest that subjects with high metacognitive efficiency accumulated more evidence after the Type 1 decisions (Fig. 11C). The post-decisional evidence accumulation allowed subjects with high metacognitive efficiency to improve the detection of correct choices (not shown in this figure), but influenced even more the detection of the incorrect choices.

It is important to emphasize that the level of certainty each rating (or wager) might represent critically depend on the difficulty level and on individual ability to accumulate and/or read out evidence. On harder trials, subjects predominantly use low wagers and, due to low average certainty (i.e. high difficulty level), these wagers represent low levels of trial certainty. On easier trials, subjects more often use high wagers because of high average certainty (i.e. low difficulty level) but, in this case, low wagers represented high or low levels of trial certainty depending on subjects’ metacognitive efficiency. Subjects with high metacognitive efficiency are able to accumulate inconsistent post-decisional evidence at high rates. This evidence not only could cross the Type 1 criterion, but also reach over far to the other side. The readouts of this evidence generate high certainty of being incorrect, which is mainly reported through the lowest wager. Subjects who do not accumulate inconsistent post-decisional evidence at high rates, on the other hand, use low wagers to report certainty readouts about the evidence that was close to the Type 1 criterion (i.e. low certainty level). This reasoning emphasizes that the same response scale can be used to represent different metacognitive readouts. Furthermore, the dependence of certainty representations on the difficulty and individual biases highlights the importance of the baseline measures for distinguishing between trial-specific and average certainty readouts.

Previous studies often interpreted low confidence reports as low certainty of being correct (Fleming and Lau, 2014; Heereman et al., 2015; Maniscalco and Lau, 2012; Sandberg et al., 2010). In such cases, low certainty readouts are expected to increase on harder trials together with the use of low ratings. However in the present study, certainty of being incorrect increased on easier trials (Fig. 10), further indicating that on these trials, subjects were using low wagers when they were more certain about their (incorrect) decisions, and not when they were more uncertain.

It is likely however that the readouts of certainty of being incorrect, or error monitoring, arise naturally in different experimental contexts, even when not specifically prompted by metacognitive report scale formulations (Yeung & Summerfield, 2012). In case of a unidirectional scale, e.g. confidence low to high, some subjects might use the middle of the scale to signal low certainty of being either correct or incorrect, and allocate low ratings for the certainty of being incorrect. Yet others might use the low ratings to designate low certainty, thus conflating these two different readouts. This issue underscores the importance of scale interpretation in addition to the formulation.

## 5. Conclusions

Comparing post-decisional judgments to a pre-decision wagering baseline allowed us to isolate individual psychological biases and expectations about task difficulty from trial-specific readouts, and to separately quantify contributions of certainty of being correct and certainty of being incorrect to metacognitive ability. Our findings demonstrate that when afforded an opportunity, humans are able to monitor and report their implicit post-decisional confidence by reading out certainty about correct and incorrect decisions. Together, readouts of a confidence in having done a correct decision, and of a certainty in the opposite direction (about having committed an error), shape metacognitive evaluations in a bi-directional manner. The error monitoring in particular drives high metacognitive efficiency. These results contribute to the ongoing discourse on complex relationship between post-decisional processing, confidence, and error monitoring. Additionally, our experimental design provides a future perspective for studying bi-directional certainty readouts not only in humans but also in other animals, as well as in patients with moderate impairments of language comprehension.

## Acknowledgements

We thank our subjects for the commitment, Lukas Schneider and Danial Arabali for important input, Dr. Holger Sennhenn-Reulen for support with statistical analysis, and members of the Cognitive Neuroscience Laboratory, German Primate Center, and the Dept. of Cognitive Neurology, UMG, for useful discussions. We also thank Dr. Stephen Fleming for his insightful and constructive comments on the manuscript, and Dr. Hakwan Lau for his valuable input during the earlier presentation on this work. This work was supported by the Brazilian Council for Scientific and Technological Development (CNPq, Science without Borders Program) grant number 201735/2012-1 to Caio M. Moreira, and the Herman and Lilly Schilling Foundation grant to Melanie Wilke.

## Supplementary material

### Archived data

The data for this article are available at the Open Science Framework, https://osf.io/ys8cj

### S.1. Supplementary Results

#### S.1.1 Wager-specific proportions of trials and slopes for the five difficulty levels

The Supplementary Figure S1A and Supplementary Figure S1B illustrate – for PreDW and PDW, respectively – the wager-specific proportions of correct and incorrect trials separately for the five difficulty levels. These proportions resulted in the slope-based measures illustrated in the third row of the Supplementary Fig. S1C and in the Figure 10. To test, within each difficulty level, if slope-correct was different from slope-incorrect depending on the trial type (PreDW or PDW), we performed two-way ANOVAs for repeated measures. In none of the difficulty levels PreDW slope-correct was significantly different from PreDW slope-incorrect (p>0.05; Supplementary Fig. S1A), indicating reliable baselines for the five difficulty levels separately.

Next, we performed paired t-tests within each difficulty level to test for significant differences between PDW and PreDW slopes. The t-tests revealed that subjects’ PDW slopes-correct were different from their PreDW slopes-correct at all difficulty levels (p<0.01), except for the highest difficulty level 5 (p>0.05). Additionally, PDW slope-incorrect was not different from PreDW slope-incorrect for the difficulty levels 4 and 5 (p>0.05), while subjects were able to read out certainty of being incorrect at the difficulty levels 1, 2 and 3 (p<0.01).

The Supplementary Figure S1 reveals two important points that are considered in our analyses. Firstly, the baseline condition (PreDW trials) showed the general (non-trial-specific) effect of expected perceptual difficulty assessments. For instance, the realization of increased family difficulty made subjects to use low wagers increasingly but independently of the correctness of the trials. Without considering this baseline measurement, we would not be able to distinguish this adaptive strategy (wagering low for hard trials) from the use of trial-specific certainty during PDW trials. Even if the task would contain only one difficulty level, or maybe especially in those cases, the baseline measurement is essential to quantify certainty readouts taking into account individual biases. Secondly, during PreDW trials (especially during expected easier trials, partially predictable by the family difficulty) subjects chose more often middle wagers (wagers 3 and 4). Since linear fits captured wagering trends regardless of the effect of such behavior, our choice for using the slopes of linear fits (instead of the best-fitted curves) proved to be valuable for establishing useful baselines for the slope-based measurements.

**Supplementary Figure S1.**
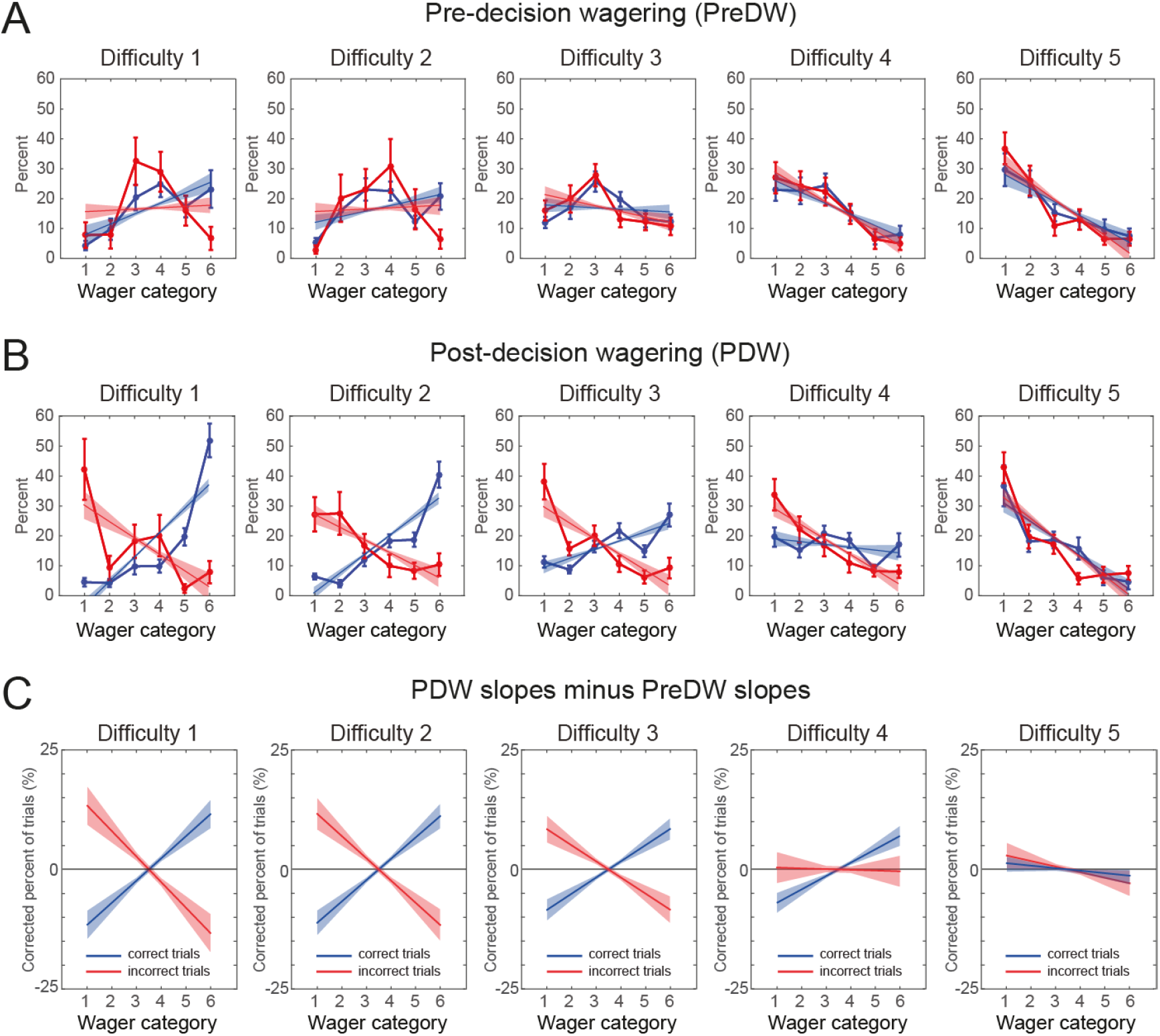
(A) Means and standard errors of linear fits for correct trials (blue lines and shaded bands) and incorrect trials (red lines and shaded bands) for (A) PreDW (baseline) and (B) PDW, fitted to the data: means and standard errors of wager-specific proportion of correct (blue curves) and incorrect (red curves) trials, for each difficulty level. (C) Means and standard errors of PDW slope-correct minus PreDW slope-correct (blue line and band) and of PDW slope-incorrect minus PreDW slope-incorrect (red line and band) for each difficulty level (same data as in Figure 10). Measurements represent averages across subjects.

The frequency of the use of each wager (means and standard errors from the lowest to the highest wager) in PDW trials (wagers 1 to 6: 19±2% 11±1% 16±1% 16±2% 13±1% 25±3%) and in PreDW trials (wagers 1 to 6: 15±2% 19±3% 23±3% 19±2% 11±2% 13±3%) showed that subjects used the entire wager scale.

#### S.1.2 The increase in Type 1 performance after wagering high in PreDW

Supplementary Figure S1A also revealed that there was an increase in Type 1 performance after wagering high in PreDW trials. For the low difficulties 1 and 2, this effect was significant for the highest wager 6, in isolation, and also reached significance after correcting for multiple comparisons across all 6 wagers for the difficulty 2 (difficulty 1: t_16_=2.45, p=0.026; difficulty 2: t_16_=3.10, p=0.0068; p=0.041, Bonferroni-corrected). We interpret this effect as a form of attentional mobilization, leading to an increase in performance after wagering high. This finding resembles a similar influence of subjective beliefs about own competency that were induced by manipulated social-comparative performance feedback and fictional research findings (Zacharopoulos et al., 2014).

#### S.1.3 Perceptual decision criteria for the five difficulty levels

We calculated the perceptual decision criterion separately for the five different difficulty levels using Signal Detection Theory approach. This information is relevant because subjects might develop different spatial biases for different difficulties (e.g. select more often the image on the right side in harder trials), making it impossible to compare different difficulty levels. The two-way ANOVA for repeated measures revealed that perceptual decision criterion did not differ from zero (F_1,16_=0.009, p=0.93) or among the difficulty levels (F_4,64_=0.248, p=0.91), suggesting that subjects identified the match on the right or left side of the screen with the same probability at all difficulty levels.

### S.2 Supplementary Discussion

Different aspects of the task design might prompt or limit post-decisional evidence accumulation and, consequently, bi-directional certainty readouts. We discuss these aspects below together with our results.

#### Time pressure

Yeung and Summerfield (2012) and others have proposed that late drifts towards the correct option, not considered by the time subjects commit to the wrong option, generate extra evidence used for error detection. In the study of Charles et al. (2013), for example, subjects who needed to report Type 1 decisions within 1 s showed higher metacognitive efficiency (meta-d′/d′, probably because the time pressure lowered d′ but not meta-d′), than those who had twice the time to report their perceptual decisions. These results indicate that time pressure over Type 1 decisions increased the use of extra information during Type 2 decisions. In our experiment, the average readout of certainty of being incorrect and the predominance of subjects with meta-d′>d′ suggest that the applied time pressure (1.5 s) was enough to restrict evidence accumulation before Type 1 decisions and favor the use of extra evidence from post-decisional accumulation or a parallel metacognitive processing.

#### Memory

memory-based tasks such as DMTS might promote the use of short-term memory as another source of extra information for the Type 2 decisions. Since the basic information required for Type 1 decisions, the previously presented sample, can only be accessed through top-down memory retrieval, such mechanism – which does not depend exclusively on continuous input of sensory evidence – might continue to provide information also after Type 1 decisions (Magnussen and Greenlee, 1999; Yu et al., 2015).

#### Propriosensory evidence

the propriosensory evidence related to the manual report of the Type 1 decision itself can serve as another source of post-decisional information. It is known that Type 1 reaction times correlate with certainty (Fetsch et al., 2014; Kiani and Shadlen, 2009). Therefore, subjects could, in principle, read out those reaction times – instead of or in addition to the perceptual evidence – in order to judge their Type 1 performance during the Type 2 decisions. In accordance with this reasoning, Fleming et al. (2015) modified subjects’ Type 2 decisions by manipulating the activity of motor areas prior to metacognitive reports. In our experiment, however, the distribution of wager-specific Type 1 reaction times (RT1) was unidirectional (decreasing towards the highest wager, Figure 9A). Even if RT1s were bi-directionally distributed, e.g. similar to metacognitive Type 2 RT2 inverted U-shape, this would only allow reading out certainty level, but not *certainty direction* (i.e. correct or incorrect). Additional possibility could be that distributions and ranges of correct and incorrect RT1s differed enough to provide a probabilistic readout of both certainty level and direction. This does not seem to be the case in our data (see Supplementary Figure S2). While the slopes fitted to incorrect wager-specific RT1s were shallower than for correct RT1s (RT1 slope-incorrect = −0.0136; RT1 slope-correct = −0.0301; t_16_=−2.148, p<0.05), their distributions overlapped substantially, especially in the low wagers. For example, in the high metacognitive efficiency group, both correct and incorrect RT1 distributions were unidirectional, with negative slopes originating from the same range in the lowest wager. These patterns render it unlikely to enable inferring the direction of certainty from a single trial RT1 ‘sample’. In summary, the readouts of RT1 in our experiment could not provide the bi-directional certainty readouts apparent in slope-based measurements and wager-specific Type 2 reaction times.

**Supplementary Figure S2.**
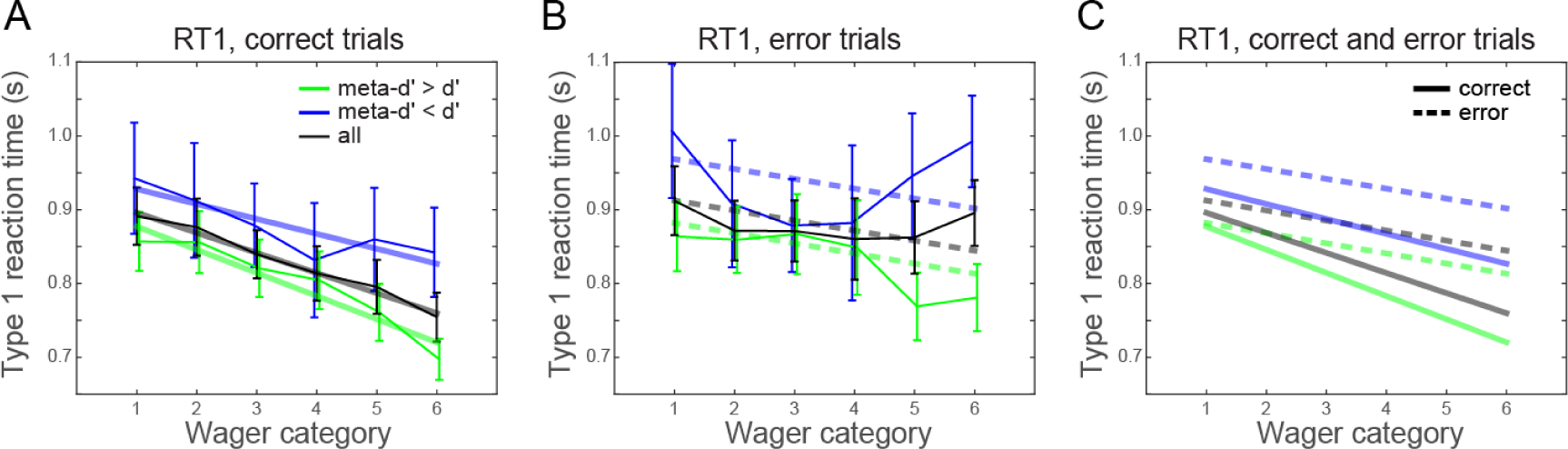
Means and standard errors of wager-specific perceptual reaction times (RT1) across all subjects (black curve), for the high metacognitive efficiency group (meta-d′>d′, green curve) and for the low metacognitive efficiency group (meta-d′<d′, blue curve), separated to correct trials (A) and incorrect trials (B). The linear regression slopes (thick lines) are means of the slopes that were fitted to the individual subject averages. Note that several subjects did not select high wagers in incorrect (error) trials; therefore data for those wagers represent only a subset of subjects while slopes include all subjects (in subjects with missing wagers, the slopes were fitted to available wagers). This led to a partial mismatch between the data and fitted slopes in B. (C) Direct comparison of correct trial (solid lines) and error trial (dashed lines) slopes.

#### Rewards and punishments

lastly, the use of PDW might favor evidence accumulation because it motivates subjects to fully explore their sources of information through gains and losses (i.e. profit more when correct and avoid losses when incorrect). We suggest that the monetary motivational aspect of PDW also contributed to the predominance of subjects who showed high metacognitive efficiency (11 out of 17 subjects).

## Abbreviations

(PreDW): pre-decision wagering
(PDW): post-decision wagering
(DMST): delayed match-to-sample task

